# Cognitive control activity is enhanced by single pulse transcranial magnetic stimulation over prefrontal and premotor areas

**DOI:** 10.1101/2023.02.02.526808

**Authors:** Jesús Cespón, Maria Concetta Pellicciari, Carlo Miniussi

**Author notes:** Corresponding author: Jesús Cespón, Institution: Basque Center on Cognition, Brain and Language, Address: Mikeletegi Pasealekua, 69, 20009 Donostia / San Sebastián, Spain, Email address.

## Abstract

Cognitive control, which includes a set of processes to implement goal directed actions and flexible behaviour, is related to a set of brain areas comprising prefrontal, premotor, and parietal cortex. However, the functional relationships between these areas and the neural correlates underlying the specific cognitive processes linked to cognitive control (e.g., inhibition and working memory update) are still unclear. In the present study, participants performed a spatial cognitive control task (i.e., a Simon task) during a transcranial magnetic stimulation electroencephalogram (TMS-EEG) co-registration. In different blocks of the task, single pulse TMS was applied over the left prefrontal, premotor, and parietal regions (a sham TMS condition was also included) at 180ms after the stimulus onset. Behavioural differences between the four TMS conditions were not observed. Accordingly, activity to inhibit the response toward the attended location was not modulated by TMS, as indexed by the contralateral central negativity (N2cc), even if TMS over parietal regions accelerated the visuospatial processing, as evidenced by faster contralateral posterior negativity (N2pc). Importantly, we observed larger P300 amplitude when delivering TMS over prefrontal and premotor cortex compared to the sham condition. These results suggest that TMS applied over the left prefrontal and premotor regions could enhances working memory processes linked to switch-and-update of the stimulus-response binding and align with the existence of prefrontal-premotor connections.

## Introduction

Cognitive control encompasses a set of cognitive processes that include working memory updating, attentional control and response inhibition. These cognitive processes allow us to implement goal directed actions (Diamond, 2013; Miyake and Friedman, 2012), which are important to carry out numerous essential daily life activities such as driving and cooking. Cognitive control processes are disrupted in numerous disorders (e.g., dementia, attention deficit and autism) (Ferguson et al., 2021; Johnson, 2012). Therefore, studying neural regions linked to cognitive control activity and how this activity could be modulated through non-invasive brain stimulation techniques such as transcranial magnetic stimulation (TMS) may be useful to implement intervening approaches that could help to improve or restore cognitive control abilities.

To investigate cognitive control, a large number of studies used the Simon task paradigm (Leuthold, 2011; Lu and Proctor, 1995). This task requires attending to a given hemifield in which a target stimulus appears and responding to a stimulus attribute (e.g., the colour) by typically pressing a lateralized response button with the corresponding hand. Even if the stimulus location is irrelevant to perform the task, the reaction time is slower and the number of errors is higher when the side of the required response is spatially incongruent with respect to the hemifield in which the stimulus is located. This interference is known as Simon effect and it has been related to the time taken to inhibit the tendency to react towards the attended location (Cespón et al., 2013; Valle-Inclán, 1996).

Several studies used TMS to test the role of specific cortical areas in the implementation of cognitive control during the performance of the Simon task. Specifically, a few studies applied TMS over prefrontal areas, as these are regions typically related to cognitive control processes (Friedman and Robbins, 2022). For instance, Stürmer et al (2007) applied repetitive TMS over the left dorsolateral prefrontal cortex (DLPFC) during the performance of a Simon task and observed altered sequential congruence effects, which suggested that TMS modulated the switch and update of the stimulus-response (S-R) binding (Cespón et al., 2020). Similarly, taking into account that dPM activity is thought to prevent the bias to respond towards the direction of the spatial attention (Leuthold and Schröter, 2006; Praamstra and Oostenveld, 2003), some studies applied TMS over dorsal premotor (dPM) cortex. Whereas some of these studies did not observe behavioural modulations (Stürmer et al., 2007), others reported increased Simon interference effect after applying single pulse TMS (Bardi et al., 2015) and repetitive TMS (Praamstra et al., 1999) over the dPM. Also, given the role of the parietal cortex in visuospatial attention (Colby and Goldberg, 1999), other studies investigated behavioural effects of applying TMS over parietal areas, showing that interference was reduced after repetitive TMS (Stürmer et al., 2007) and even suppressed in a study using a single pulse TMS protocol (Schiff et al., 2011).

A limitation of previous TMS studies is the absence of cortical measures to investigate how TMS modulates neural correlates of cognitive processes that are crucial to perform the Simon task such as deployment of spatial attention to the target stimulus, activity to monitor and inhibit the response towards the direction of the spatial attention, and working memory updating of the stimulus-response binding (Cespón et al., 2020). By analysing correlates of neural activity, it would be possible to identify neural modulations underlying behavioural changes induced by TMS.

Given its high temporal resolution, event-related potentials (ERP) is a suitable tool to study how TMS modulates visuospatial attention and cognitive control during the performance of the Simon task. There are specific ERP correlates related to attention to a target location and inhibition of the tendency to react towards the attended location. Specifically, the contralateral posterior negativity (N2pc) is a correlate of visuospatial attention to the target stimulus and suppression of the non-target stimulus (Luck, 2012; Mazza et al., 2009; Woodman and Luck, 2003; Zivony et al., 2018), which arises from posterior-occipito-temporal areas (Hopf et al., 2000) and the contralateral central negativity (N2cc) is an ERP correlate of neural activity to monitor and prevent the tendency to react towards the attended location (Cespón et al., 2012; Praamstra and Oostenveld, 2003) -which emerges from the dPM (Oostenveld et al., 2001; Praamstra, 2006). There are additional ERPs that have been studied during the performance of cognitive control tasks; namely, the fronto-central N200 and the P300 (Cespón and Carreiras, 2020). Fronto-central N200 was related to conflict monitoring activity emerging from the anterior cingulate cortex and the amplitude of this component is usually increased at higher task demands (Folstein and Van Petten, 2008). Also, P300 is a correlate of switching and update of the S-R binding (Adrover-Roig and Barceló, 2010; Hoppe et al., 2017), whose neural sources were mainly located in parietal and cingulate cortex (Frühholz et al., 2011; Linden et al., 2005) and its amplitude directly correlates with enhanced working memory skills (Polich, 2007).

In the present study, we applied single pulse TMS at 180ms after the onset of the stimulus over F3, FC5, and CP5 electrodes -which target the left DLFPC, dPM, and supramarginal/angular regions, respectively (Koessler et al., 2009) - in order to investigate behavioural and neural modulations of cognitive control processes during the performance of a Simon task. The different areas were stimulated in different blocks of the task (see Figure 1). A sham TMS condition, with the coil located over the vertex (Cz), was also used. TMS was applied only over the left hemisphere due to its key role in visuomotor transformations (Rushworth et al., 2001; Spironelli et al., 2006).

**Figure 1.**
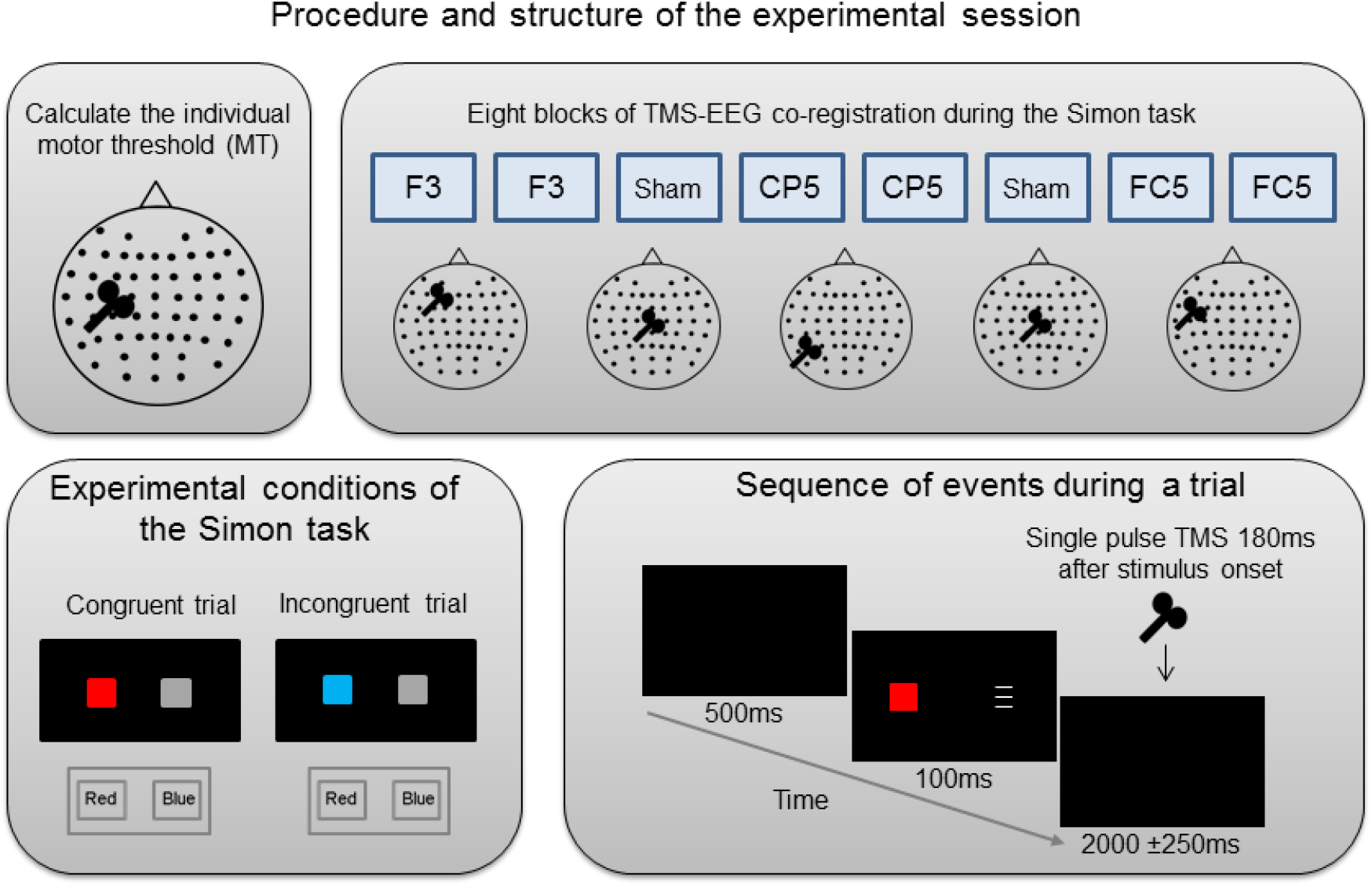
Procedure of the experimental session and Simon task. The top part of the figure graphically summarizes the procedure of the experimental session: electrodes montage, motor threshold (MT) calculation and performance of the different blocks of the Simon task during TMS-EEG co-registration. The bottom part of the figure represents the two experimental conditions (congruent and incongruent) and the sequence of events during a trial.

We expected that delivering a single TMS pulse would alter on-going neural activity and interfere with performance (Miniussi et al., 2013). So, we hypothesized that TMS over supramarginal/angular gyrus would slow reaction times in the congruent and incongruent conditions by delaying the time to allocate attention to the target stimulus, which would be revealed by slowed N2pc latency. Even so, TMS over supramarginal/angular region would not change the Simon effect, unless the dPM activity is also altered, which may occur by modulating the activity throughout a parietal-premotor (visuomotor) pathway and such changes could be indexed by N2cc. Considering the closed left DLFPC-left dPM connections (Abe and Hanakawa, 2009), we predicted that TMS over these two areas would increase the interference (i.e., Simon effect) by slowing the neural processes reflected by N2cc and P300. Finally, we expected that fronto-central N200 would be larger in the incongruent than congruent conditions and it could also be increased in those TMS blocks with augmented interference effect.

## Methods

### Participants

Fifteen young adults (11 females, 4 males; mean age = 23.2 SD = 2.39) took part in the study after being screened to exclude any participant with contraindications for TMS (Rossi et al., 2009). Participants were right-handed, as assessed by the Edinburgh handedness inventory test (Oldfield, 1971), and had normal or corrected to normal visual acuity. Participants reported no previous history of neurological and/or psychiatric disorders. Experimental protocols for TMS were performed in accordance with the suggested checklist for a routine clinical TMS examination recommended by Rossini et al. (2015). This research was conducted in compliance with the ethical guidelines agreed by the Declaration of Helsinki and received prior approval from the local Ethics Committee. Before taking part in the experiment, all participants were informed about the procedures of the study and provided written informed consent.

### Procedures

The whole experiment was conducted in a dimly lit and shielded room. At the beginning of the experimental session, the individual motor threshold (MT) was calculated as the intensity in which a motor evoked potential (MEP) of 50μV was observed in the electromyogram (montage in first dorsal interosseous of the right hand) the 50% of the times for at least 10 TMS pulses delivered over the left primary motor cortex through a standard 70 mm figure-of-eight coil (Magstim Super Rapid Whitland, UK). The calculation of the MT was carried out after placing the EEG cap, which ensured to deliver the right intensity given the spacing created by the electrodes mounted in the cap. After the MT calculation, participants performed eight blocks of the Simon task paradigm (see below or Figure 1) during the EEG recording for about one hour. On each subject, two blocks of TMS were delivered over F3, FC5, and CP5, which targeted left DLPFC, dPM and angular/supramarginal gyrus, respectively, according to estimations by Koessler et al. (2009). In addition, the participants received two blocks of sham TMS over Cz (sham TMS), in which 3 cm spacer was placed between the coil and the scalp. TMS pulses were delivered 180ms after the onset of the stimulus at an intensity of 80% of the MT. The structure of the session is represented in the Figure 1. The order of the stimulated area was counterbalanced among the participants. During the stimulation, the coil was placed tangentially on the scalp, with the handle pointing backward at an angle of 45° from the mid-sagittal axis of the participant’s head. Subjects wore protective headphones to minimize the auditory response produced by the TMS coil click. A neuronavigation system (SofTaxic Optic EMS, Bologna, Italy) permitted a constant control of the stability and spatial consistency of the coil position and its orientation throughout the stimulation period.

### Task

During the Simon task, participants, who were sat 100 cm in front of the screen, were asked to direct their gaze to the centre of the display. A grey fixation cross appeared in the centre of the screen for 500ms against a black background. Then, a bilateral array of stimuli appeared for 100ms and participants had to respond as fast and accurate as possible to the colour of the stimulus (a red or blue square) by pressing one of the two response buttons arranged horizontally. A target stimulus (i.e., the square) and a non-target stimulus (a grey contralateral stimulus with similar shape, size, and lateralized location) were presented 5 cm to the left or right of the central fixation cross so that the entire display was within the foveal region (i.e., around 3 degrees of visual angle from the centre of the foveal region) (Bargh and Chartrand, 2000). The presence of a contralateral non-target stimulus avoids any influence from exogenous asymmetries in the EEG while it does not alter the spatial Stimulus-Response compatibility effects (O’Leary and Barber, 1993; Leuthold, 2011). After the bilateral presentation of the stimuli, the screen remained blank for a time period of 2000 ±250ms and then a new trial began with the appearance of the central fixation cross. The task consisted of 120 trials per block (so, for every condition, there were 120 x 2 = 240 trials; 120 congruent and 120 incongruent trials). Considering that there were eight blocks, the total number of trials in the task was 960 (480 congruent and 480 incongruent trials, presented in a random order). Before starting the task, participants performed a practice block of 10 trials. The task is graphically represented in Figure 1.

### EEG recording

TMS-compatible EEG equipment (BrainAmp 32MRplus, BrainProducts GmbH, Munich, Germany) was used to record the EEG from 57 sintered, Ag/AgCl passive electrodes (EasyCap, Brain Products GmbH, Germany). The ground electrode was placed at Fpz. The right mastoid served as an online reference for all electrodes, whereas the left mastoid electrode was used offline to re-reference the scalp recordings to the average of the left and the right mastoid, i.e., including the implicit reference (right mastoid) in the calculation of the new reference. The EEG signal was acquired with a bandpass filter of 0.01-1000 Hz and digitized at a sampling rate of 5000 Hz (for conducting the analyses reported in this article, the sampling rate was down-sampling to 1000 Hz). Vertical and horizontal electrooculogram signals were recorded by two electrodes located above and beneath the right eye and two electrodes located lateral to the outer canthi of both eyes. The impedance between the skin and EEG electrodes was below 5 kΩ.

### Data analysis

Behavioural performance was evaluated by analysing reaction times (RTs) and accuracy (Number of Errors – NE). ERP analyses included N2cc, N2pc, N200, and P300. EEG data were linearly interpolated between −1 to 19 ms from each TMS pulse (i.e., between 179 and 199 ms after the onset of the stimulus). The EEG signal was filtered with a 0.1-80 Hz digital bandpass and a 50 Hz notch filter. Ocular and muscular artefacts were corrected by using independent component analysis [algorithm Infomax (Gradient) applied to the whole dataset implemented in Brain Vision Analyzer]. Epochs showing activity higher than ±100 μV were automatically rejected. All remaining epochs were individually inspected and removed from subsequent analyses if residual artefacts were still detected. In order to analyses event-related potential modulations induced by TMS pulses, epochs were extracted from −200 to 800 ms with respect to target stimulus presentation. A baseline correction (−200 to 0 ms) was carried out.

ERPs were calculated only for the correct responses. Following previous studies (Cespón et al., 2020), the N2pc was obtained based on the hemifield of the target stimulus location by applying the following formula: [P8 - P7 (left hemifield stimuli) + P7 - P8 (right hemifield stimuli)] / 2]. N2cc was obtained applying the same formula to C3 and C4 electrodes. For N2pc and N2cc components, we did not distinguish between congruent and incongruent conditions in order to increase the signal/noise ratio and to avoid an overlap with residual motor activity in central areas (Praamstra, 2007). The N2cc and N2pc peak latencies were identified as the largest negative peaks between 200–300ms after stimulus presentation. Also, for each subject in congruent and incongruent conditions separately, we created the following two regions of interest (ROI) to study N200 and P300 components: fronto-central ROI (by pooling the electrodes Fz, FCz, FC1, FC2, and Cz) and parietal ROI (by pooling the electrodes CPz, Pz, P1, P2, and POz). Based on the grand averages, N200 peak latency was identified as the maximum negative peak between 240ms and 320ms after the onset of the stimulus and P300 was identified as the maximum positive peak between 280ms and 420ms after the onset of the stimulus. N200 and P300 amplitudes were calculated in time windows of 20ms and 50ms, respectively, around their peak latencies.

### Statistical analysis

To analyse the behavioural data (i.e., RT and number of errors -NE), we used a repeated measures ANOVA with Condition (two levels: Congruent, Incongruent) and Stimulation (four levels: CP5, FC5, F3, and sham) as within-subject factors. Also, given that TMS was applied in different areas over the left hemisphere, we wanted to exclude a different TMS effect on the basis of the hemifield in which the target stimulus was located. For this purpose, the mentioned repeated measures ANOVAs for RT and NE (that is, Condition x Stimulation) were carried out by using Hemifield of the target stimulus (two levels, Left target stimulus, Right target stimulus) as an additional within-subject factor.

To analyse N200 and P300 latencies and amplitudes, we carried out the corresponding repeated measures ANOVAs with Condition (two levels: Congruent, Incongruent), Stimulated site (four levels: F3, FC5, CP5, Cz), and ROI (two levels: fronto-central and parietal) as within-subject factors. For N2pc and N2cc latencies and amplitudes, we implemented a repeated measures ANOVA using Stimulated site (four levels: F3, FC5, CP5, Cz) as within-subject factor.

When any of conducted ANOVAs showed significant effects related to the main factors or their interactions, the Bonferroni correction was applied to post hoc tests. Also, if the condition of sphericity was not met, then the Greenhouse-Geisser correction for degrees of freedom was applied. Finally, we reported the partial eta square (η2p) as measure of effect size for the significant results.

## Results

### Behavioural results

The repeated measures ANOVAs including Hemifield as a within-subject factor (that is, Condition x Stimulation x Hemifield) did not produce any significant effect related to Hemifield. The repeated measures ANOVA for RT (Condition x Stimulation) showed a significant effect of Condition [F (1, 14) = 29.9, p < 0.001; η^2^p = 0.682], as the RT was slower in the incongruent than congruent condition (p < 0.001). The Stimulation and Stimulation x Condition interaction did not produce significant effects.

Also, the repeated measures ANOVA for NE revealed a significant effect of Condition [F (1, 14) = 12.4, p = 0.003; η2p = 0.471], as the NE was higher in the incongruent than congruent condition (p = 0.002). The Stimulation and Stimulation x Condition interaction did not show significant effects. Slower RT and higher NE in incongruent compared to congruent conditions are graphically represented in Figure 2.

**Figure 2.**
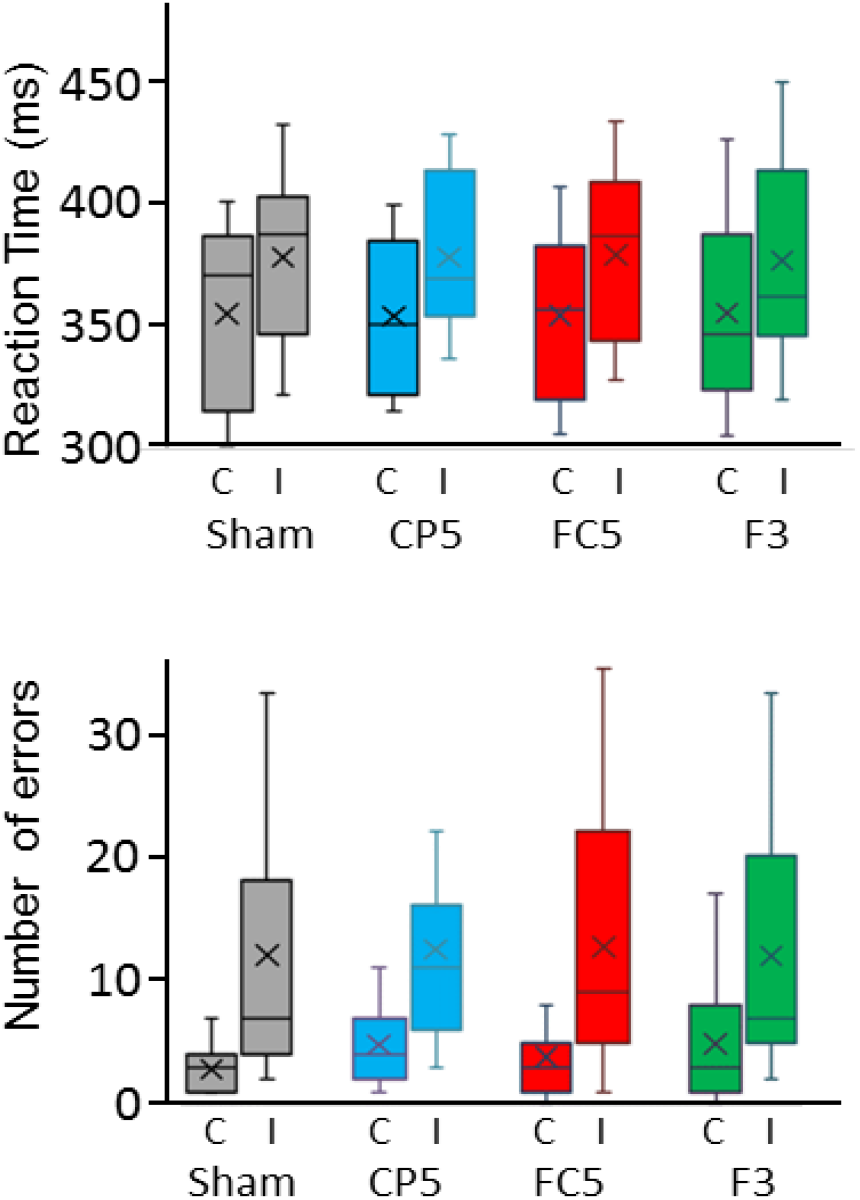
Representation of the behavioural results. Reaction times (RT) and number of errors (NE) for congruent (C) and incongruent (I) conditions in the blocks with sham TMS over Cz and real TMS over CP5, FC5, and F3. Spatially incongruent trials showed slower RTs and higher NE compared to congruent trials.

### Event-related potentials (ERP)

The repeated measures ANOVA for N200 amplitude (Figure 3) revealed an effect of the condition [F (1, 14) = 8.59, p = 0.011; η^2^p = 0.380], as N200 was larger in incongruent than congruent condition. Also, an effect of the ROI was observed [F (1, 14) = 46.26, p < 0.001; η2p = 0.768], as N200 amplitude was larger in fronto-central than in parietal ROI. For P300 amplitude, the repeated measures ANOVA (Stimulation x Condition x ROI) showed an effect of Stimulation [F (2.2, 30.8) = 5.82, p = 0.002; η2p = 0.294], as P300 amplitude was larger when stimulating over F3 than sham (p = 0.037) and over FC5 than sham (p = 0.041) (see Figure 3). No significant effects were observed for N200 and P300 latencies.

**Figure 3.**
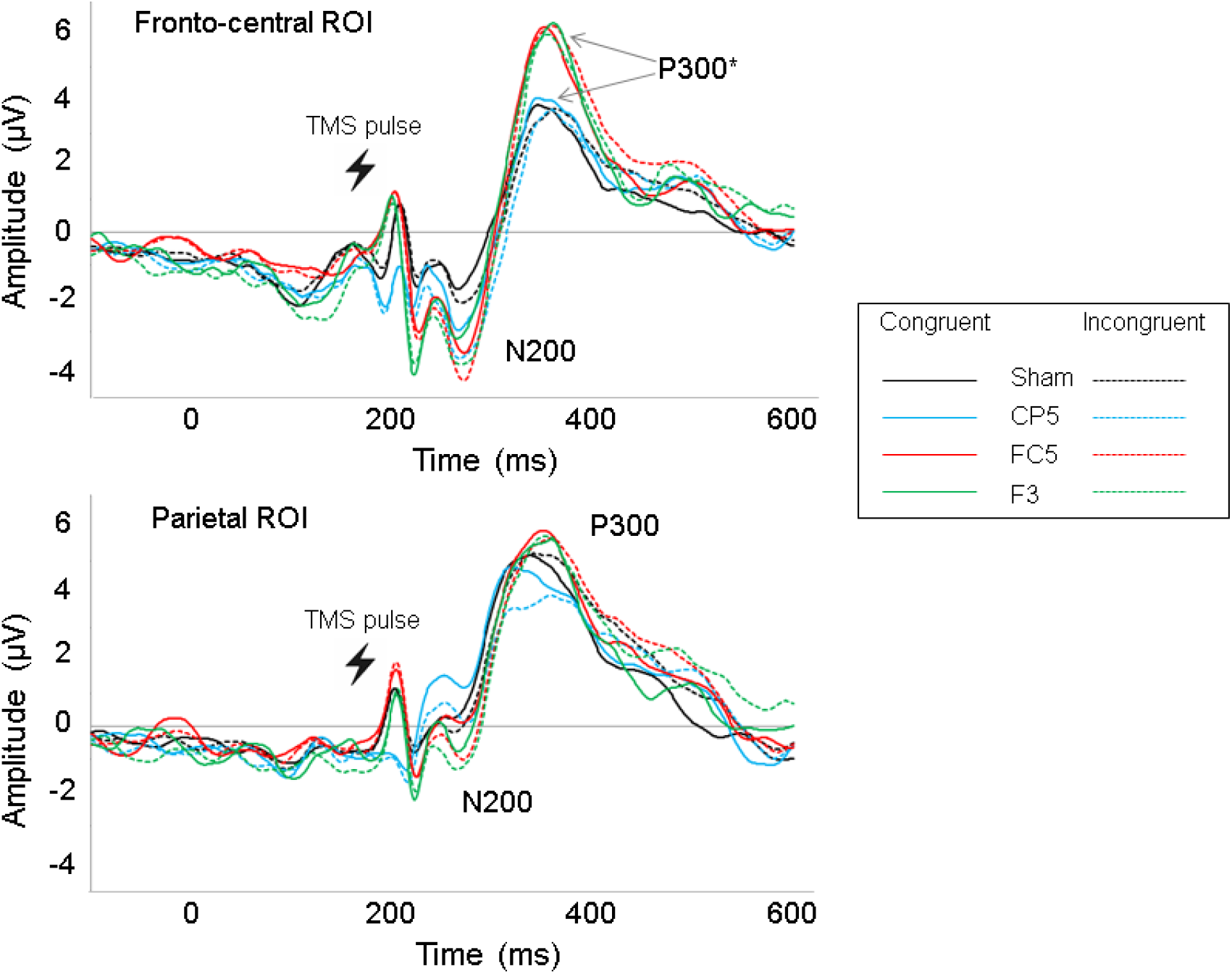
N200 and P300 in fronto-central and parietal regions of interest (ROIs). In fronto-central ROI, P300 amplitude was larger when applying TMS over FC5 and F3 compared to the other conditions (sham over Cz and CP5). In addition, fronto-central N200 amplitude was larger in incongruent than congruent trials.

The repeated measures ANOVA for N2pc latency (Stimulation) revealed a significant effect [F (3, 42) = 5.80, p = 0.002, η2p = 0.293], as the N2pc latency was faster in those blocks in which TMS was delivered over CP5 compared to F3 (p = 0.013) and sham (p = 0.043) conditions (see Figure 4). The analysis for N2pc amplitude did not reveal any significant effect. The repeated measures ANOVAs for N2cc latency and amplitude did not reveal any significant effect.

**Figure 4.**
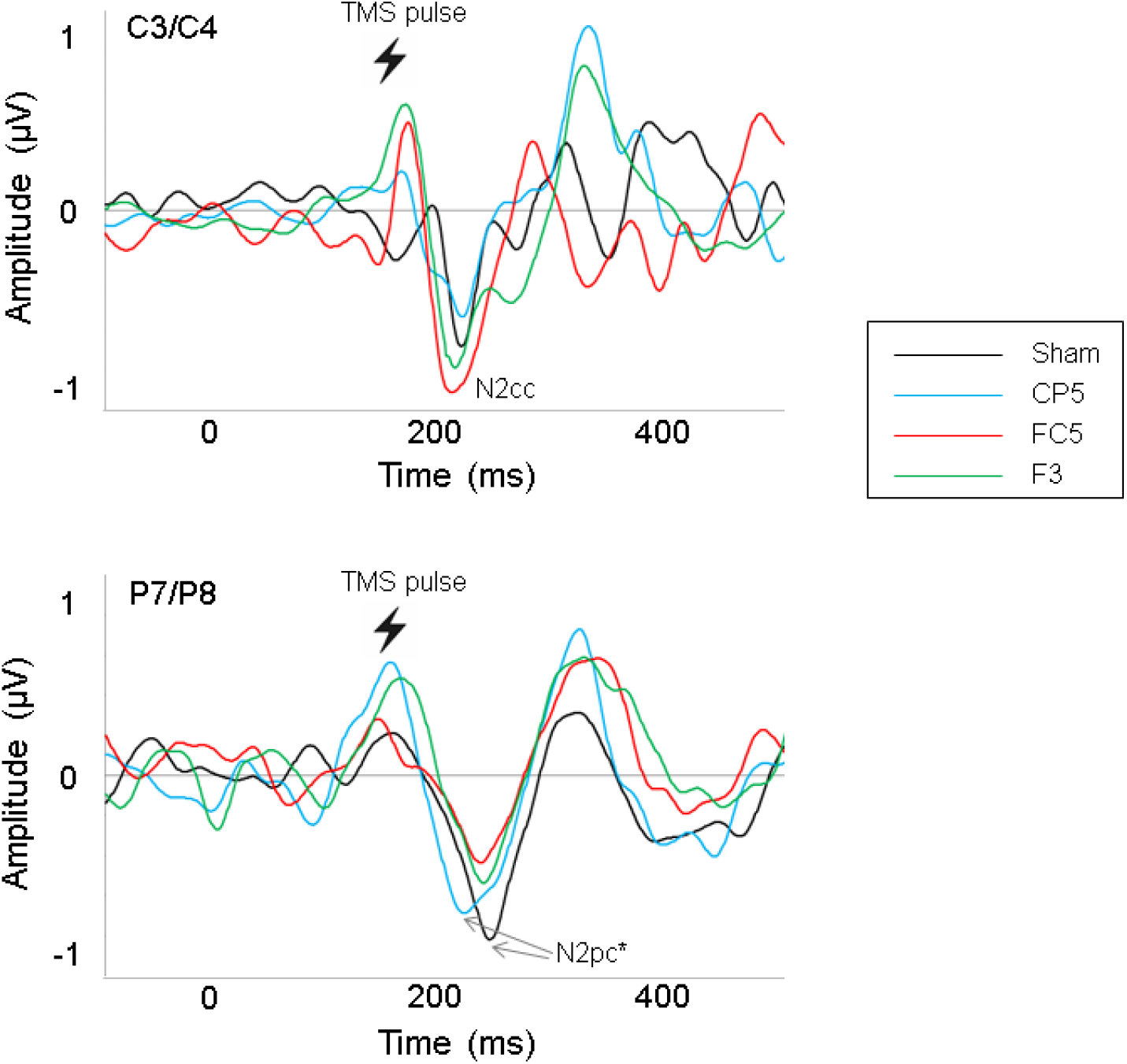
Contralateral central negativity (N2cc) and contralateral posterior negativity (N2pc). N2pc was analysed in P7/P8 electrodes pair because it is where N2pc achieved the maximum amplitude. N2pc latency was significantly earlier when applying TMS over CP5 compared to the blocks of sham TMS and TMS over F3.

## Discussion

The results of the present study showed slower reaction times (RT) and higher number of errors (NE) in the spatially incongruent compared to congruent conditions of the Simon task. Also, fronto-central N200 amplitude was larger in incongruent than congruent conditions. The TMS did not produce different behavioural modulations -as revealed by RT and NE-on the basis of the stimulated site (i.e., F3, FC5, CP5, and sham TMS over Cz). However, the amplitude of P300 was larger when TMS was delivered over F3 and FC5 compared to the blocks in which TMS was applied over CP5 and sham TMS. Moreover, N2pc latency was faster with TMS delivered over CP5 compared to F3 and sham TMS.

The behavioural results of the present research showed the well-established Simon effect, as revealed by increased RT and NE in incongruent compared to congruent conditions (Cespón et al., 2020; Lu and Proctor, 1995). Also, ERPs showed increased fronto-central N200 amplitude in incongruent than congruent conditions, which suggests augmented cognitive control activity from fronto-central midline regions and anterior cingulate cortex in order to cope with conflicting information (Folstein and Van Petten, 2008). Increased fronto-central N200 in conflicting than non-conflicting trials is consistent with previous research using diverse cognitive control paradigms (Cespón and Carreiras, 2020; Folstein and Van Petten, 2008). Instead, P300, which was also investigated in both congruent and incongruent conditions separately, was not modulated by the Simon effect. The absence of P300 modulations related to the Simon effect is consistent with the view of P300 as an ERP component that is mainly modulated by switching the S-R binding rather than by the Simon effect itself (Cespón et al., 2020; Hoppe et al., 2017); that is, P300 amplitude is usually lower in those conditions in which participants have to switch the S-R binding from the previous trial, which happens in congruent trials preceded by incongruent trials as well as in incongruent trials preceded by a congruent trials.

The main objective of the present study was investigating the behavioural and neural modulations related to TMS pulses. In contrast to our predictions, we did not observe any behavioural effect related to TMS. In any case, these are not highly surprising results, as previous studies applying TMS during the performance of the Simon task reported facilitation, interference as well as no behavioural effects (Cespón et al., 2020). As discussed in the following paragraphs, the obtained electrophysiological results align with a neural facilitation effect of the TMS; specifically, increased P300 amplitude by delivering TMS over F3 and FC5 and faster N2pc latency by delivering TMS over CP5.

The TMS delivered over F3 (i.e., left DLFPC) and FC5 (i.e., left dPM) increased the P300 amplitude. Therefore, in line with our predictions, TMS over prefrontal and premotor cortex similarly modulated ERP correlates of cognitive control. It is possible that premotor-prefrontal connections (Abe and Hanakawa, 2009; Schulz et al., 2019) explain that TMS over both regions similarly modulate the P300 component. According to previous studies, increased P300 amplitude relates to enhanced cognitive functioning as well as lower effort to perform the task (Amin et al., 2015; Polich, 2007; Russo et al., 2008). So, in the context of the Simon task performance, the results of the present study suggest that TMS over F3 and FC5 augmented neural activity to update the S-R binding (Cespón et al., 2020; Hoppe et al., 2017) even if such signal change was no reflected at the behavioural level. These findings are in line with improved executive functioning after low frequency repetitive TMS over the frontal lobe (Bashir et al., 2020) with the difference that in the present experiment, there is only a single stimulus.

On the other hand, TMS over FC5 or F3 did not produce any effect on N2cc, a correlate of dPM activity to monitor and inhibit the tendency to react towards the attended location (Cespón et al., 2020; Praamstra and Oostenveld, 2003). It is possible that TMS over FC5 modulated N2cc only in the incongruent condition (in which inhibitory activity is deployed to prevent the tendency to react towards the attended location) but not in the congruent condition, in which such inhibitory activity is not needed. Future studies using a higher number of trials would be able to test this hypothesis by distinguishing between congruent and incongruent conditions of the task and applying specific methods –e.g., residual iteration decomposition (Ouyang et al., 2015)- to disentangle activity related to N2cc from motor activity, as both sources of activity overlap in central regions (Praamstra, 2007).

The N2pc latency had an earlier peak when stimulating over CP5 -which was intended to stimulate angular and supramarginal parietal areas-compared to sham and F3 TMS. These results might point to a facilitation effect to allocate attention to the target stimulus and/or inhibit the non-target stimulus, which represent the cognitive processes reflected by N2pc (Mazza et al., 2009; Woodman and Luck, 2003; Zivony et al., 2018). Nevertheless, such facilitation revealed by earlier N2pc peak did not give rise to improved performance and/or earlier sending of motor outputs to premotor cortex to implement cognitive control of the tendency to react towards the hemifield of the stimulus location, as N2cc was not modulated by TMS over CP5. A hypothetical explanation for these results is that earlier N2pc peak latency after TMS over CP5 was driving by processes linked to non-target suppression rather than target processing. For this reason, earlier N2pc peak latency does not necessarily involve earlier N2cc peak latency (i.e., faster deployment of neural activity to prevent a reaction towards the attended location).

We have pointed to the relatively low number of epochs as a limitation that make no possible to distinguish between congruent and incongruent conditions for N2pc and N2cc components. Another limitation of the present study that should be highlighted is that we reported TMS effects when TMS was delivered at 180ms after the stimulus onset. However, other studies have observed behavioural effects using other timings such as TMS targeting the left dPM (Praamstra et al., 1999) and the left DLPFC (Stürmer et al., 2007) several hundred of milliseconds before the onset of the stimulus. A more comprehensive research could include a first experimental session to assess the timing in which TMS effectively modulates the performance and, subsequently, using TMS-EEG co-registration to investigate the neural underpinnings of the observed behavioural modulations.

Another issue is that TMS clicks could have induced an “accessory effect”, which speeds up the cognitive processing (Jepma et al., 2009; Tona et al., 2016). Thus, it is unclear whether and to what extent the increased levels of arousal related to the “accessory stimulus” could have masked or attenuated differences caused by the different TMS conditions. In order to shed light on this issue, future research could use a control block delivering sham TMS (as we did in the present study) and, additionally, another block in which participants perform the task without the use of any (real or sham) TMS.

In summary, single pulse TMS applied over the left prefrontal, premotor and parietal regions during the performance of a cognitive control task (specifically, a Simon task) did not produce any significant behavioural modulation compared to the sham TMS condition. In contrast, we observed several electrophysiological modulations induced by TMS. Namely, earlier N2pc peak latency, a correlate of visuospatial attention to the target stimulus and inhibition of the non-target, was observed after delivering TMS over parietal areas. However, such modulation was not accompanied by faster reaction times or earlier sending of a motor command to precentral regions, probably because earlier N2pc peak latency was driven by modulated non-target suppression rather than target processing. Importantly, the main finding of the present study was that TMS over the left prefrontal and premotor areas enhances the P300 amplitude, a correlate of working memory updating, during the Simon task performance. Thus, the results of the present study suggest the utility of applying single pulse TMS over prefrontal and premotor regions to strength neural processes related to working memory update.

## Acknowledgements

This study was funded by European Commission Marie-Skłodowska Curie Actions, Individual Fellowships; 655423-NIBSAD, from the Basque Government through the BERC 2018-2021 program, from the Agencia Estatal de Investigación through BCBL’s Severo Ochoa excellence award SEV-2015-0490 and by the CARITRO Foundation.

JC: designed the study, collected the data, analysed the data, and wrote the manuscript.

MCP: supported data analyses and provided a critical revision of the manuscript.

CM: designed the study and provided a critical revision of the manuscript.

